# Mechanistic constraints on the trade-off between photosynthesis and respiration in response to warming

**DOI:** 10.1101/2023.06.29.546917

**Authors:** Suzana G. Leles, Naomi M. Levine

## Abstract

Phytoplankton are responsible for half of all oxygen production and drive the ocean carbon cycle. Metabolic theory predicts that increasing global temperatures will cause phytoplankton to become more heterotrophic and smaller. Here we uncover the metabolic trade-offs between cellular space, energy, and stress management driving phytoplankton thermal acclimation and how these might be overcome through evolutionary adaptation. We show that the observed relationships between traits such as chlorophyll, lipid content, C:N and size can be predicted based on the metabolic demands of the cell, the thermal dependency of transporters, and changes in membrane lipids. We suggest that many of the observed relationships are not fixed physiological constraints but rather can be altered through adaptation. For example, the evolution of lipid metabolism can favor larger cells with higher lipid content to mitigate oxidative stress. These results have implications for rates of carbon sequestration and export in a warmer ocean.

**Teaser:** A tale of how photosynthetic microbes might defy current trends to become larger and grow faster in a warmer ocean.

## Introduction

Ocean temperatures are increasing at unprecedented rates due to anthropogenic increases in carbon dioxide in the atmosphere (*1*). These rising temperatures have the potential to substantially alter both rates of carbon cycling in the ocean and marine ecosystem dynamics (*2*–*4*). Here we focus on the response of phytoplankton to increasing temperatures as they drive carbon cycling in the ocean, contribute half of all oxygen production, and form the base of the marine food web (*5, 6*).

Phytoplankton have a classic unimodal response of growth rate to temperature (*7*). This response can be broken down into several competing processes. As temperatures increase, cellular metabolism speeds up as enzyme kinetics increase due to more collisions between the substrate and the active site of the enzyme (*8, 9*). This results in faster rates of carbon fixation and respiration (*10*). In addition, warmer temperatures increase rates of protein denaturation (*9*). Overall, respiration rates have been shown to increase more rapidly than carbon fixation with increasing temperature (*10*). However, the specific metabolic constraints driving this relationship is unclear.

Understanding the fundamental molecular and energetic trade-offs that underlie classic trait relationships is critical for predicting how phytoplankton will adapt to a warming ocean. Conventionally, predicted responses have been based on present day trait correlations from empirical evidence. For example, that warming decreases body size (*3, 11*) and drives ecosystems to become more heterotrophic (*2, 6*). However, we lack a fundamental understanding of the molecular mechanisms behind these two hypothesized responses and thus an understanding of how robust future predictions are on the timescales of anthropologically driven climate change.

Here we ask whether observed correlations between temperature and phenotypic traits can be altered while still maintaining fundamental molecular, energetic and physical constraints. This is motivated by recent work which showed that some trait correlations (e.g. between metabolism and size) can be altered through adaptation (*12*). In fact, an experimental evolution study found that phytoplankton evolved larger cell sizes and higher growth rates due to warming (*13*), opposed to the expected smaller cell sizes (*14*).

By leveraging tools from thermodynamics, resource allocation theory and trait ecology, we developed a proteome model for phytoplankton to investigate the mechanistic constraints on photosynthesis and respiration due to warming. The model optimizes proteome allocation and cell size to maximize phytoplankton growth under different temperatures. We use this model to provide new mechanistic insights into the trade-offs between respiration, cellular space and environmental stress management that limit the metabolic choices of phytoplankton acclimation under temperature stress and how these might be overcome through adaptive change. Our results suggest that phytoplankton acclimate to warming by switching the main respiratory pathway from glycolysis to lipid degradation due to a trade-off based on proteomic cost, energy yield and space constraints. In addition, decreases in cell size are driven by the thermal dependency of nutrient uptake and by the increased saturation of membrane lipids. We suggest that the relationship between size and temperature can be overcome through changes in lipid metabolism (e.g. increasing lipid content to mitigate oxidative stress). Thus, under certain environmental conditions, phytoplankton might adapt to warming by evolving larger cell sizes with faster growth rates. This is consistent with experimental data that found that the most rapidly evolving genes were associated with redox homeostasis, oxidative stress and transcriptional regulation (*13*), all functions that could be tightly linked to lipid metabolism. Understanding the hard and soft constraints that drive phytoplankton acclimation and adaptation will allow us to better predict their impact in the carbon cycling and trophic efficiency as the climate changes.

## Results

### A proteome model for a phytoplankton cell

An illustrative schematic of the single-cell proteome allocation model for phytoplankton thermal responses is shown in Figure 1A. We use the model to solve for the daytime proteome, macromolecular concentrations, and surface area to volume ratio of a photosynthetic cell as a function of temperature. We assume that the cell grows under steady-state ideal light and nutrient conditions. While we do not simulate diurnal cycles, the cell must generate sufficient protein pools and stored carbon to fuel respiration during the night. Resources are allocated to different protein pools (shown in blue), which regulate the production and consumption of other macromolecules (shown in green).

**Figure 1.**
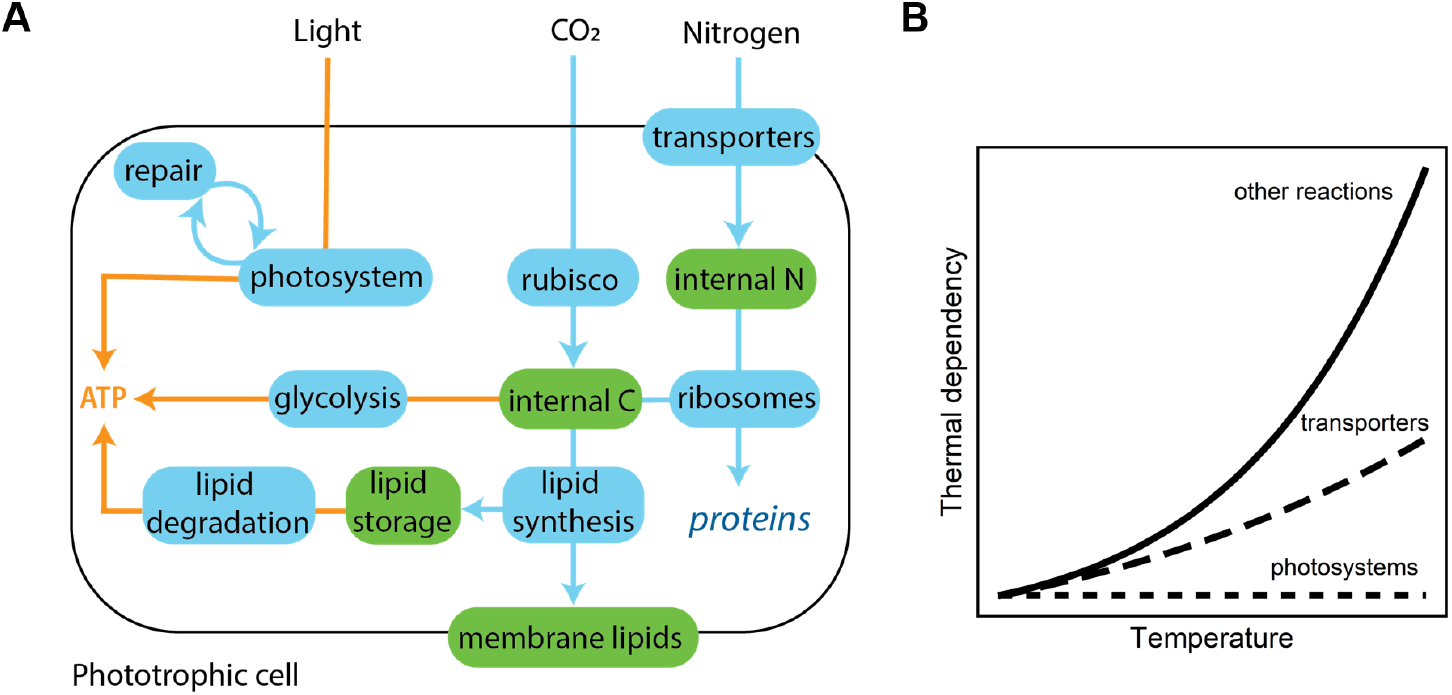
The single-cell proteome model for phytoplankton. (A) Blue pools indicate proteins and green pools indicate other macromolecules. Arrows indicate one-way reactions with blue arrows indicating the production of a given macromolecule catalysed by a given protein and orange arrows indicating energy fluxes. The model optimizes the relative investment in the different proteins. (B) The model assumes that photosystems are temperature independent and that the thermal dependency of nutrient transporters is lower than that of other reactions (see Section “A proteome model for a phytoplankton cell”).

Photosystems convert light energy into chemical energy. The conversion of inorganic carbon into a generic internal carbon pool (akin to carbohydrates) is mediated by Rubisco activity. Nitrogen transporters located in the cell membrane bring inorganic nitrogen into the cell. The internal pools of carbon and nitrogen are used by ribosomes to synthesize new proteins, including the ribosomes themselves. The internal carbon pool is also used for lipid synthesis and to fuel dark respiration through glycolysis. The lipid synthesis pathway is used both to build membrane lipids and for producing stored lipids. Stored lipids can be used to fuel dark respiration (*15*) through the lipid degradation pathway. Because our model optimizes for the proteome of the cell in a steady-state environment, glycolysis and lipid degradation are mutually exclusive; the optimal cell exclusively uses the more efficient pathway given a specific temperature. In the model, stored lipids can also be used to minimize heat damage (Eqs. 21 and 22). The synthesis and accumulation of triacylglycerol molecules (TAGs) into lipid droplets has been shown to be a protective mechanism through which both phytoplankton and plants cope with stress conditions (*16, 17*). Lipid droplets help maintain energy and redox homeostasis mitigating oxidative stress and also regulate membrane composition by removing unsaturated lipids at elevated temperatures (*17*–*19*).

In order to optimize for the surface area to volume ratio of the cell, the model accounts for a trade-off between having more membrane real estate for nitrogen transporters and having more biovolume for the proteome and carbon storage. The surface area of the cell is equal to the area required for the nitrogen transporters and the membrane lipids that are required for membrane integrity (Eqs. 37-39). Both carbohydrates and lipid storage take up cellular space and thus require larger biovolumes (i.e. smaller surface area to volume ratios). However, these two carbon storage pools have different impacts on cell size because lipids are hydrophobic and form lipid droplets that do not impact osmoregulation (*15*). Therefore, we assume that stored lipids, while contributing to cell volume, do not contribute to a maximum allowed density (Eqs. 41-43). The model can consider cells of different shapes but here we assumed a spherical cell for simplicity.

Warming accelerates all enzymatic reactions in the model, except for photochemistry (Figure 1B) because photon capture is temperature independent (*20*). However, Rubisco and thus carbon fixation are temperature dependent, thus photochemistry and carbon fixation are modeled separately. In addition, the model can impose different thermal dependencies for different reactions (e.g. Figure 1B).

The model also accounts for the fact that the lipid per unit area of membrane increases with temperature (Eq. 27). This results from the switch from unsaturated to saturated lipids at warmer temperatures (*21*). Since saturated lipids pack more easily in the membrane due to the lack of kinked hydrocarbon chains, the cell requires a greater concentration of membrane lipids to maintain membrane integrity at higher temperatures.

Finally, the model assumes that protein damage increases with increasing temperature (Eqs. 19 and 20). Although intracellular damage due to elevated temperatures is likely to affect all proteins, photosystems are particularly vulnerable because heat damage is commonly accompained by light stress (*18, 22*). Thus, we assume warming results in increased damage to photosystems and thus requires repair by specialized proteins (Eqs. 23 and 24), following Mairet *et al*. (*23*). A version of the model where all protein pools are damaged at high temperatures and require repair is provided in the Supplement; this model showed qualitatively similar results. The model and parameters are fully defined in the supplement (Tables S1-S3).

### Model comparison against experimental data

Our proteome optimization model of a phytoplankton cell reproduces observed trait values and trends as a function of temperature (Figures 2 and S1). Here we compare the model output against diatom physiological measurements. We selected diatoms due to the abundance of trait data and the availability of experimental evolution studies for this group. By accounting for the acceleration of enzymatic reactions alongside photosystem damage as a function of temperature, our model captures observed growth and carbon fixation rates of various diatom species as well as the observed decrease in the carbon use efficiency with warming (Figure 2A,B and C). In agreement with empirical data, the model also predicts an increase in the chlorophyll content per unit of biovolume, a decline in the nitrogen to carbon ratio of the cell, and a decrease in cell size as temperature increases (Figures 2D, E and F). These responses are independent of the respiratory pathway used by the cell (Figures 2 and S1). The model then allows us to tease apart the underlying molecular mechanisms driving these commonly reported relationships between traits and temperature.

**Figure 2.**
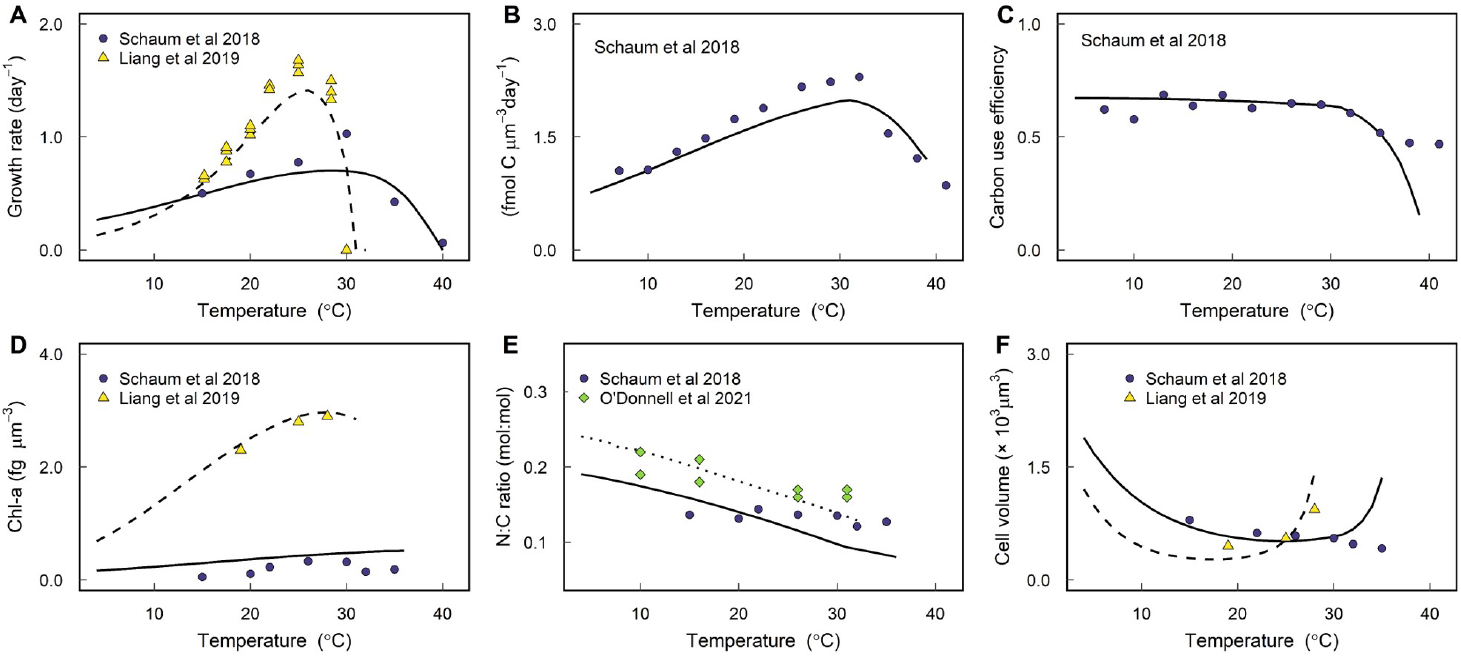
Model validation for thermal acclimation. Comparison between model (lines) and experimental data (symbols) is shown. Experimental data are for various diatom species acclimated to different temperatures. Solid lines correspond to model calibrated to data from Schaum *et al*. (*13*), dashed lines to Liang *et al*. (*24*) and dotted line to O’Donnell *et al*. (*14*) (Table S3). Here we assume the cell could only invest in glycolysis to fuel dark respiration but model validation was also performed assuming that the cell could only invest in lipid degradation (Figure S1).

### Shifts in investment with increasing temperatures

Differential temperature dependencies of cellular processes play a critical role in observed trait relationships as the cell acclimates to warming. If temperature had the same effect on all cellular processes, no shift in allocation would occur, as increased energy generation would keep pace with increased respiration demands. This result would be inconsistent with observations. In contrast, decoupling temperature independent photon absorption by photosystems from the temperature dependent enzyme kinetics of Rubisco results in an energetic imbalance. Thus, as temperature increases, the optimal cell invests in more light-harvesting proteins and divests resources away from Rubisco in order to maximize growth rates (Figures 3A and S2A). This generates the observed increase in chlorophyll-a per biovolume as a function of temperature (Figure 2D). Photosystem damage due to temperature stress exacerbates this imbalance as the cell must also invest in more repair proteins as temperature increases (Figures 3B and S2B) to avoid the excessive accumulation of damaged photosystems (Figure S3). Our main results do not change if all proteins are assumed to be damaged at the same rate as the photosystems at high temperatures, although this modified version of the model incurs a higher penalty in growth rates as temperatures increase due to the greater investment in repair proteins and respiration (Figure S4).

**Figure 3.**
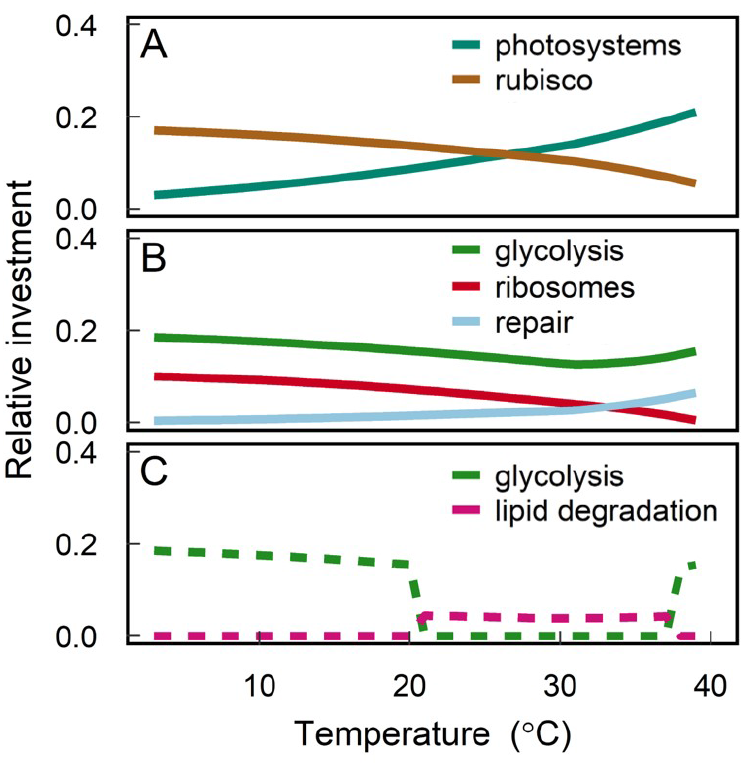
Optimal proteome investments. Changes in the relative proteome investment in different protein pools as a function of acclimated temperature. Solid lines (panels A and B) correspond to the investments when glycolysis is the only respiratory pathway available to the cell (investments are qualitatively the same when lipid degradation is the only respiratory pathway available; Figure S2). Metabolic switches are observed when the cell is given the option to choose between glycolysis or lipid degradation to fuel respiration (panel C). Model output is shown for the model calibrated to data from Schaum *et al*. (*13*) and fitted values are given in Table S3.

The optimal cell compensates for this need to increase investment in photosystems and repair proteins by divesting resources away from other parts of the proteome. Increased kinetic energy results in more efficient pathways for protein synthesis, thus the optimal cell can decrease the relative investment in ribosomes without impacting growth rates (Figures 3B and S2B). Thus the model predicts a decrease in the concentration of ribosomes with increasing temperatures (Figure S5), providing a mechanism for the observed decrease in ribosome concentration with warming (*25*). The increased investment in photosystems, and resulting decrease in investment in ribosomes, contributes to the observed decrease in the nitrogen to carbon ratio of the cell with increasing temperatures (Figures 2E), as photosystems are less nitrogen rich than ribosomes (Table S2).

As temperature increases, the optimal cell initially divests resources away from respiratory pathways to accommodate the increased photosystem demand. This pattern is ultimately reversed as temperatures increase to near fatal temperatures due to exponentially increasing metabolic demands (Figures 3B and S2B), which we discuss more below. If the cell is given a choice between fueling dark respiration with glycolysis or lipid degradation, the optimal cell switches from using glycolysis to using lipid degradation at a critical high temperature (Figure 3C). This result is consistent with transcriptomic and proteomic evidence from cultured isolates, in which glycolysis was found to be up-regulated in colder environments (*26*) while lipid degradation was up-regulated in warmer environments (*24*). Here we suggest that this switch arises due to a trade-off between energy demands, protein requirements, and space. To meet increasing energy demands due to rising temperatures, a cell must balance the relative investments in uptake and growth. While it costs less energy to build carbohydrates, lipids store more energy per mole of carbon (*27*) and also take up less space in the cell (*15*). The model predicts that lipids are preferentially used for respiration at higher temperatures in order to support higher metabolic demands while optimizing for smaller cell sizes (Figure S6), which helps to maintain a high surface area to volume ratio (see discussion below). By using lipid degradation instead of glycolysis, the cell can also increase its carbon use efficiency and the relative concentration of photosystems in the cell (Figure S6). Once temperature increases to a near fatal temperature, the optimal cell shifts into a third regime. In this regime, both respiration demand and stress due to photosystem damage are excessively high. Under these conditions, the optimal cell stores lipids to regulate stress rather than to fuel respiration and switches back to glycolysis to meet respiration demands (Figure 3C). Consequently, the required carbohydrate storage to fuel dark respiration results in larger optimal cell sizes (Figure S6). A cell can avoid entering into this third regime by increasing the efficiency of the lipid degradation pathway (Figure S7). A short coming of the proteome allocation model is that it optimizes assuming a steady-state environment. As a result, the model never predicts the utilization of both the lipid degradation and glycolysis pathways, resulting in the metabolic switch (Figures 3C and S6). In reality, we expect that fluctuating environmental conditions would generate a smooth transition in which both pathways are utilized over the transition range.

### Changes in cell size due to warming

Our model allows us to explore the impact of having different thermal dependencies for different reactions. We found that the thermal dependency for nitrogen uptake relative to other reactions 1B) plays a significant role in the observed changes in cell size with warming (Figure 2F). Specifically, if the thermal dependency of nutrient uptake is equal to that of other metabolic processes (e.g. ribosomes), as temperatures increase, the cell can maximize growth rate by divesting resources away from transporters in order to invest in the synthesis and repair of photosystems. This results in a decrease in the relative concentration of transporters per unit of biovolume as temperatures increase (Figure S8A), resulting in a smaller surface area to volume ratio and larger cell sizes (Figure 4A). This is opposite to the observed trends of decreasing cell size with increasing temperatures even under replete nutrient conditions (*11*). On the other hand, if we assume that nitrogen transport into the cell is less sensitive to temperature than other metabolic processes, nitrogen uptake is less efficient than other pathways as temperatures increase and thus the cell must increase the relative concentration of transporters to maintain the same flux of nitrogen into the cell (Figure S8A). This results in larger surface area to volume ratios and smaller cell sizes (Figure 4A).

**Figure 4.**
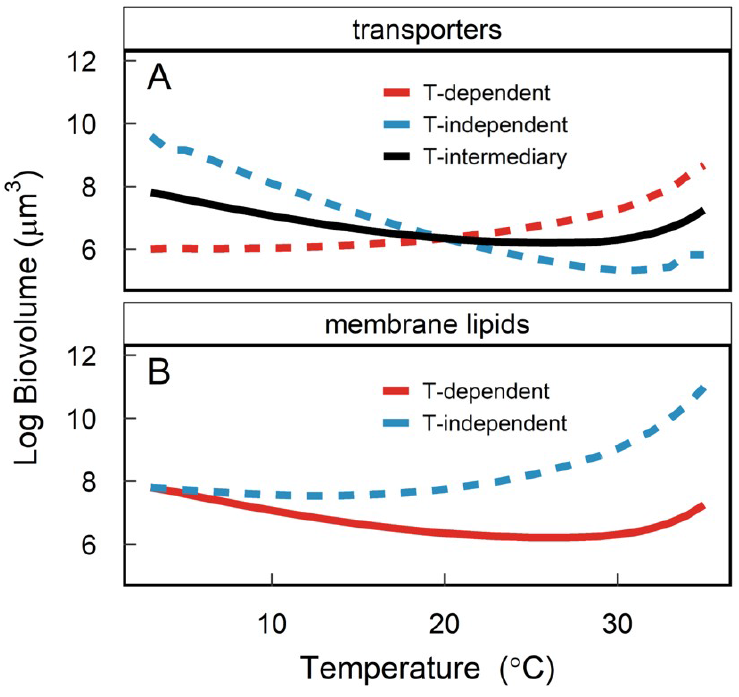
Cell size depends on the thermal dependencies of membrane components. Changes in cellular biovolume depend on the thermal dependencies of nitrogen transporters (A) and membrane lipids (B). Solid lines indicate the baseline run (shown in Figures 2 and 3) while dashed lines indicate the sensitivity runs. In panel A, the T-dependent run (red dashed) uses the same temperature sensitivity for transporters as other reactions. To provide the other extreme case, runs in which transporters are assumed to be independent of temperature are shown (blue dashed). The baseline uses an intermediary scenario for nutrient transporters (see Figure 1B). Panel B uses the intermediate temperature sensitivity for transporters and shows the scenario for which the assumption that lipid saturation changes with temperature was removed from the model. Model output is shown for the model calibrated to data from Schaum *et al*. (*13*) and fitted values are given in Table S3.

Previous experimental evidence demonstrated that the thermal dependency of nutrient uptake is lower than that of both carbon fixation and growth (*28, 29*). In addition, isotope incorporation studies and enzyme assays showed that the thermal dependency of nitrate uptake by the diatom *Thalassiosira pseudonana*matched the thermal dependency of nitrate reductase activity, which was lower than that of growth (*30*). Specifically, the *Q*_10_ (the factor by which metabolic rates increase by a 10 ^*o*^C rise in temperature) for nitrate reductase activity was found to be 1.8 (*31*) while the *Q*_10_ for Rubisco and respiration is approximately 2.4 and 3.0, respectively (*20*). Here we halved the activation energy *E*_*a*_ (Eq. 3) of transporters relative to other processes, but the model results are qualitatively the same with different decreases in *E*_*a*_ (Figure S9).

Cell size is also determined by the concentration of membrane lipids. If we assume that there is no change in lipid saturation as temperature increases, then the cell must synthesize sufficient membrane lipids to meet the membrane integrity constraint (Figure S8B), which implies that the ratio between transporters and lipids in the membrane should be lower or equal than a constant value (Eq. 26). This results in lower concentration of membrane lipids per unit of biovolume and thus smaller surface area to volume ratios and larger cell sizes (Figure 4B). However, when we incorporate the need for cells to switch from unsaturated to saturated lipids for membrane stability, the relative concentration of lipids per unit area of membrane increases (Figure S8B). Saturated lipids take up less space compared to unsaturated lipids because they lack kinked hydrocarbon chains (*15*). Therefore, to maintain a constant spacing between transporters in the membrane, the ratio of lipids to transporters must increase with increasing temperature (Eq. 27). When this change to saturated lipids at high temperatures is included in the model (Eqs. 26 and 27), the surface area to volume ratio of the cell increases, resulting in smaller cell sizes (Figure 4B).

It is noteworthy that the model predicts that cells get larger again at near fatal temperatures across all modelled scenarios (Figures 2F and 4). This occurs during the third regime described above. Energetic constraints at high temperatures force the cell to heavily invest in respiration and carbon storage, decreasing the concentration of transporters and membrane lipids per unit of biovolume (Figure S8). Consequently, the cell decreases its surface area to volume ratio to accommodate the required carbon storage and energy investment.

### Drivers of thermal adaptation

Cells respond to shifts in environmental conditions first by acclimating (rearrangement of metabolism through phenotypic changes) and then by adapting (genetic changes). The proteome model allows us to study both responses. The acclimation response is captured by the different allocation strategies predicted at different temperatures (i.e. those described above) based on a set of constant parameter values (Table S2). However, by altering underlying parameters (Table S3), we can also use the model to understand the adaptive responses of phytoplankton. The differences in the validation runs fit to the three different datasets in Figure 2 (*13, 14, 24*) is an example of different adaptations captured by the model. Here we extend this analysis and use the proteome model to investigate the potential trade-offs associated with high temperature adaptive responses and associated mechanisms driving these trade-offs. Specifically, we allow changes (akin to mutations) in parameters (akin to genes) associated with the thermal dependency and maximum turnover rate of reactions as well as the maximum lipid content and the cost associated with heat-damage (Table S3). We compare our results against data from Schaum *et al*. (*13*), in which diatom strains were evolved at different temperatures: 22 ^*o*^C (control), 26 ^*o*^C (moderate warming) and 32 ^*o*^C (severe warming).

Cells can adapt to warmer temperatures by evolving enzymes that function better at high temperatures (*32*). In the model, this is represented as a shift in the thermal dependency of the reactions (increasing the activation energy *E*_*a*_; Table S3) to allow for faster kinetics with more stable enzymes at higher temperatures. This adaptation allows higher growth rates at high temperatures, but comes at the cost of reduced growth rates at lower temperatures due to slower kinetics (Figure 5A). In addition to this known trade-off, the model predicts additional trade-offs associated with evolving enzymes that perform better at high temperatures. Since the thermal dependency of nutrient transport into the cell is lower than other reactions in the model (see discussion above), warm adapted cells can divest resources away from transporters at cooler temperatures, resulting in larger cell sizes compared to their ancestors (Figure 5B). Furthermore, because metabolic rates still increase exponentially with temperature in the warm adapted cells, the optimal cell must also store even more carbon than the ancestor cell to fuel respiration as temperatures increase (Figure 5C). Thus, at high temperature, the warm adapted cells both grow faster and are larger than the ancestral strains (Figure 5B, blue versus yellow).

**Figure 5.**
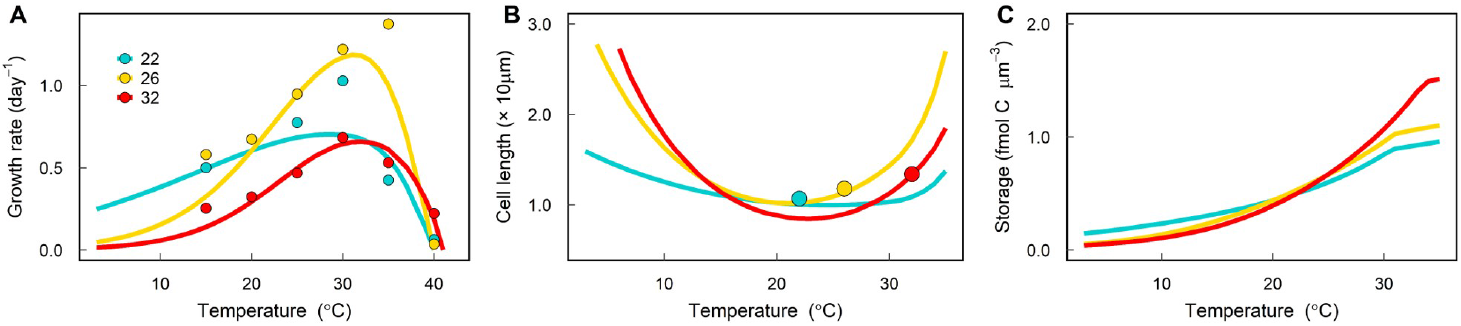
Thermal acclimation of phytoplankton evolved at different temperatures. Model (lines) comparison against experimental data (symbols). Experimental data from Schaum *et al*. (*13*) include observations for different diatom lineages evolved at three temperatures: 22^*o*^C (control), 26^*o*^C (moderate warming) and 32^*o*^C (severe warming). In panel C, carbon storage accounts for both carbohydrates and lipids required to meet dark respiration costs as well as stored lipids to regulate oxidative stress.

In order to adapt to critically high temperatures (i.e. grow faster and increase carbon use efficiency), cells must invest in mechanisms to minimize and repair damage resulting from oxidative and heat stress (*13*). While there are many ways in which a cell can invest resources to mitigate heat and oxidative stress, here we test the role that lipids might play in allowing for high temperature adaptations. In the model, stored lipids can mitigate heat damage (Eqs. 21 and 22). In addition, in the evolved runs, we allow cells to increase their maximum lipid content 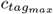 which acts to decrease the cost of heat-damaged photosystems *e*_*d*_ (Table S3). This allows cells to increase growth at extreme temperatures. The trade-off of this strategy is that it requires increased cell sizes to accommodate higher lipid storage (Figure 5). The model dynamics are consistent with experimental data which showed that cells evolved to high temperatures are larger (*13*). Phytoplankton have also been found to increase their fatty acid content relative to their ancestors after adapting to warming (*33*).

The model is able to capture the trade-offs associated with adaptation to both moderate and extreme changes in temperature. By increasing the maximum lipid content (*ctag*_*max*_), the 32°C evolved model is able to sustain growth at the previously fatal temperature of 40°C (Figure S10A). This growth at extreme temperatures comes at the cost of reduced growth across the growth curve relative to the 26°C evolved model (e.g. maximum growth of 1.02 day^1^ compared to 1.14 day^1^ for the 26°C evolved model). By evolving both higher lipid content and shifting the thermal dependency of the reactions (higher *E*_*a*_), the cell can obtain higher growth rates relative to the simulations where only maximum lipid content increased, albeit still slightly lower than the 26°C evolved model (Figure S10A). In addition, this extreme temperature strategy requires larger cell sizes (Figure S10B). These trade-offs are a result of higher investment in the synthesis of stored lipids to minimize damage, which take up cellular space and divests carbon away from growth (Figure S10C). These model results are consistent with experimental results which found that diatoms evolved under extreme temperature stress (32°C) had growth rates that were lower for all temperatures except 40°C relative to the ancestral strains or moderate temperature evolved strains (Figure 5A). While the model qualitatively captures the observed shifts in growth rates for the extreme temperature evolved strains by only considering changes in lipid content and *E*_*a*_, the model is only able to capture the larger decrease in growth rates across the entire growth curve by also decreasing the maximum turnover rates of all reactions (i.e., *k*_*ref*_, Table S3). Decreases in *k*_*ref*_ for high temperature adapted strains are consistent with observations of slower rates of photosynthesis and respiration from experimental evolution studies (*10, 13, 34*). The model suggests that a benefit to slowing all reaction rates is that it keeps energy demands lower at high temperatures, decreasing respiration rates and thus requiring less carbon storage which allows the cell to maintain smaller cell sizes (Figure S10B). More work is needed to understand the trade-offs associated with adaptations to these extreme temperatures but the model provides a useful tool for teasing mechanisms apart.

## Discussion

Our proteome model offers a unique approach for uncovering the underlying cellular mechanisms behind observed plastic and evolved phytoplankton trait changes to shifts in temperature (Figure 6). Identifying which trade-offs are hard constraints determined by energetic or biophysical limitations versus maleable constraints that can be overcome via evolution provides new insights into how phytoplankton might adapt and impact ecosystem processes in a warmer ocean (*13, 35*). We make three important predictions: i) lipids are preferentially used for respiration as temperature increases because they have a higher recoverable energy and take up less space in the cell; ii) changes in the saturation of membrane lipids and the lower thermal dependency of nutrient uptake explain why cells acclimate to warming by decreasing size and iii) phytoplankton might break the observed temperature-size correlation and evolve larger sizes under temperature stress. We hypothesize that increasing cell size will be an important strategy to increase lipid content and minimize oxidative stress when faced with critically high temperatures.

**Figure 6.**
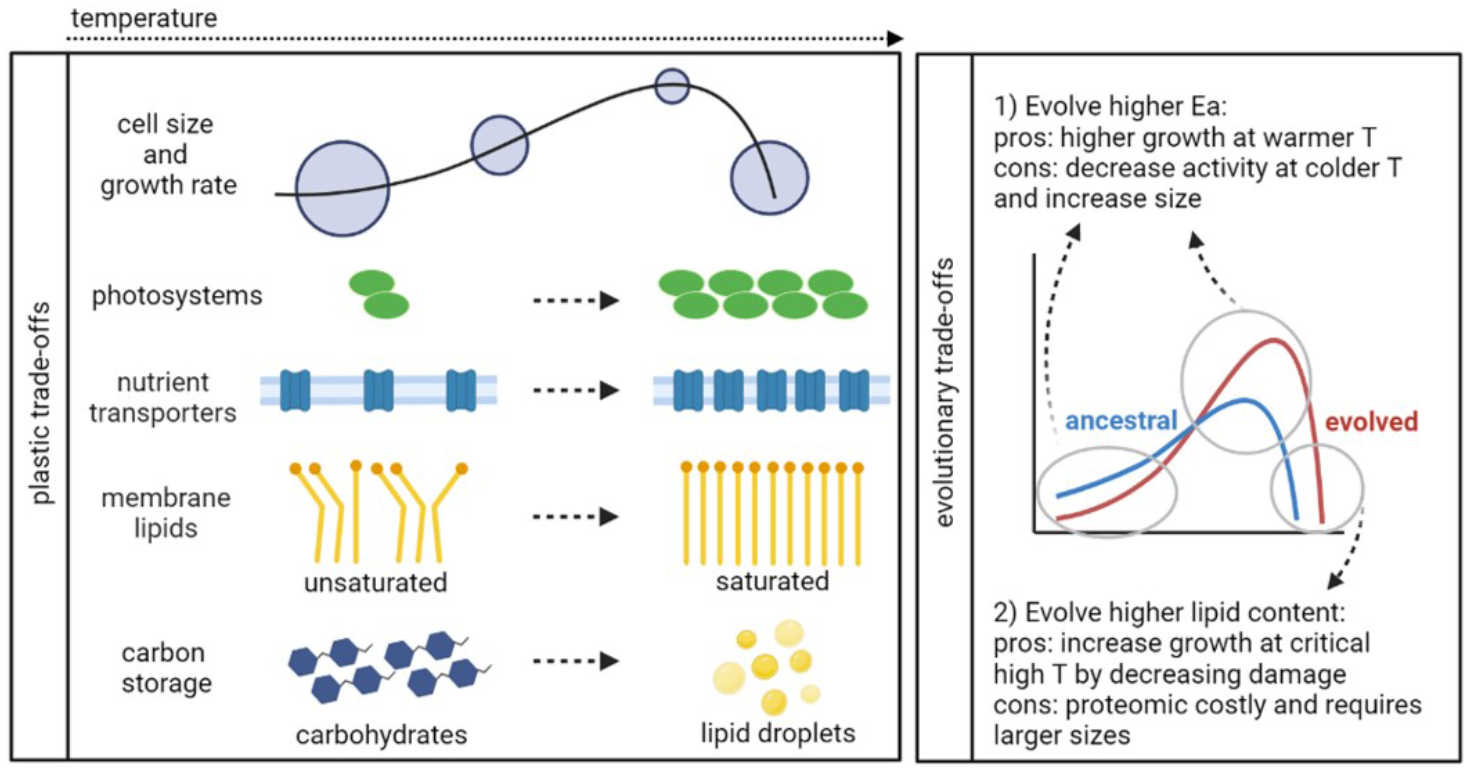
Phytoplankton thermal trade-offs. Phytoplankton respond plastically to increases in temperature by decreasing cell size, increasing photosystems, transporters and the saturation of membrane lipids, and by storing lipids instead of carbohydrates. At critical high temperatures the cell gets bigger again because it must invest all it can in carbon storage to fuel high respiration rates. To adapt to warming, phytoplankton can evolve higher *E*_*a*_ so that enzymes function better at high temperatures at the cost of higher carbon storage and lower growth at colder temperatures. They could also evolve higher lipid content to decrease cellular damage which requires larger sizes.

### Why do phytoplankton acclimate to warming by decreasing cell size?

As predicted by theory and empirical evidence, we find that phytoplankton acclimate to warming by decreasing cell size (Figure 6). Decreases in size as a function of temperature have been reported for diverse organisms, both across and within species; in fact, this functional response is so ubiquitous that it has been considered a universal ecological response to global warming (*3*), though this is debated (*36*). Decreases in the size of marine plankton communities due to warming are explained by the accompanied effects on decreased nutrient availability due to increased stratification of the water column (*4, 37*). However, decreases in cell size are also observed experimentally when warming is decoupled from nutrient limitation (*11*). Our model reveals that this is because even under replete nutrient conditions, phytoplankton can become limited by nutrients due to the lower thermal dependency of nutrient uptake relative to other metabolic processes (*20, 28, 30*). Thus, as temperatures warm, a cell must increase the relative investment in nutrient transporters to keep pace with increased metabolic demands. In addition, at higher temperatures, cells switch from unsaturated to saturated lipids (*21*), requiring a higher relative investment in lipids to maintain membrane integrity, which also decreases the optimal cell size.

We identify a trade-off between allocating cellular space to photosystems and carbon storage (increasing cellular volume) versus membrane components (increasing cellular surface area) as temperatures increase (Figure 6). To manage this trade-off, we found that the cell switches from glycolysis to lipid degradation due to warming, as supported by empirical evidence (*24, 26*). By choosing to store lipids instead of carbohydrates (*15*), phytoplankton fine tunes respiration (*38*) by more efficiently storing energy per mole of carbon as well as per unit of biovolume. This increases the cells carbon use efficiency and allows for smaller cell sizes at warmer temperatures. However, the model also predicts that cells should get larger again at critically high temperatures (Figure 6). This has been observed empirically (*24, 39*) and is usually attributed to cell cycle arrest just before cell death (*18*). Here we propose that there is an additional mechanism at play. Cells must invest more in repair and respiration, and thus carbon storage, divesting resources away from membrane components and investing resources in components that require larger cellular volume. Since lipids can help mitigate damage due to oxidative stress, our model predicts that the optimal cell switches back to glycolysis to fuel respiration, which promotes further increases in cell size at extremely high temperatures (Figure S6F).

### Can phytoplankton adapt to warming without shrinking?

One might ask: is the observed relationship between small size and warmer temperatures strongly constrained or is there a way for phytoplankton to adapt to warming without shrinking? In fact, Schaum *et al*. (*13*) found that while high temperature acclimated cells decreased cell size, high temperature adapted populations evolved larger cell sizes and higher growth rates at elevated temperatures compared to the ancestral population. Using the proteome allocation model, we reveal the mechanisms and trade-offs both behind the observed trend of decreases in cell size with temperature and how phytoplankton might alternatively be able to get larger under temperature stress, thus challenging the temperature size rule. We found that in order to evolve higher growth rates at warmer temperatures, phytoplankton can (i) store more carbon to fuel higher rates of respiration and/or (ii) decrease heat-damage by evolving higher lipid content. Both strategies require allocating more cellular space towards storage, which requires larger cell sizes. The predictions described above have important consequences to ecosystem functioning. Higher fatty acid content translates into better food quality for predators, which would support more productive food webs in a warmer ocean. Such changes are critical because they suggest that some phytoplankton might adapt to warming by getting larger (larger carbon sink) and supporting higher trophic transfer, contrary to short-term predictions (*33, 40*).

Other trait changes not considered in our modeling study could also favor larger cells. Marshall *et al*. (*12*) demonstrate that microbes can evolve to be larger, more metabolically efficient, and to have faster growth rates. Their results imply that larger cells have lower energetic costs per unit of biovolume relative to small cells. This may be due to the fact that changes in membrane lipid saturation are energetically costly (*41*). Since smaller cells have larger surface area to volume ratios, their costs to maintain membrane fluidity should be relatively higher than for larger cells. Together with larger storage capacity, these mechanisms help to explain why a large cell size could emerge as an alternative adaptive strategy due to warming, especially in environments with frequent and high variations in temperature and pulses of nutrients. In such an environment, larger cells would be able to store more carbon to regulate oxidative stress to support higher growth during warmer periods. Under the multi-stressor environment of high temperature and nutrient limitation, previous work has suggested that phytoplankton thermal responses are more strongly constrained (*42*); and thus that thermal adaptation was not possible (*43*). However, even the most oligotrophic seas are influenced by nutrient pulses derived from meso- and sub-mesoscale features (*44*). Thus, large phytoplankton could benefit from storing nutrients during these episodes, suggesting that there might be a niche for being large in a future warmer ocean.

### Future directions

Our findings are grounded on a large body of theory (*2, 45*) and lessons from trait ecology (*11, 46*) as well as on new insights from experimental evolution (*13, 33*) and omic approaches (*21, 24, 38*). By integrating these fields, we investigated the gaps in knowledge related to the molecular trade-offs that constrain both short-term acclimation and evolutionary adaptation of phytoplankton due to warming. Previous work has shown that proteome models can provide insight into microbial choices across different environments (*23, 47*–*51*). Here we show how proteome allocation models can be used to investigate short-term (acclimative) and long-term (adaptive) trade-offs. Future studies designed to test the hypotheses and predictions raised here against proteomic data are critical, first focusing on culturable isolates (*18, 26*) and then against data obtained from natural communities. Ultimately, proteome models can inform the key trait trade-offs that should be included in large-scale ecosystem models (*25, 52*), allowing us to better understand how fine-scale processes scale up to influence global biogeochemical cycles. This is particularly critical because both trait values and the correlation between traits will constrain the possible evolutionary outcomes of phytoplankton in a warmer ocean (*53*).

## Materials and Methods

### The proteome allocation problem

Our coarse-grained model is built on a constrained optimization problem that maximizes the steady-state growth rate of a generic photosynthetic cell considering temperature, irradiance levels, and external dissolved inorganic nitrogen and carbon concentrations. The model is based on previous proteome models developed for heterotrophic and phototrophic metabolisms (*23, 47*–*51*). For each environmental condition, the model optimizes the following state variables: the intracellular concentration of proteins (*p*) and other macromolecules (*c*), the relative investments in the different proteins (*β*), the volume to surface area ratio of the cell (ϕ), and the fractions of the lipid synthesis flux designated to build membrane lipids (*∝*_*lm*_), stored lipids (*∝*_*tag*_), and to fuel the lipid degradation pathway (*∝*_*ld*_) (Table S1). Our work extends previous models to account for temperature effects, lipid and energy metabolisms, and space constraints in the model, allowing us to predict changes in phytoplankton cell size and stoichiometry as a function of temperature. Parameter values are presented in Table S2 for those that were kept constant throughout our simulations and in Table S3 for those that were fitted to observational data. More details on the choice of parameter values are given in the Supplementary Methods (Tables S4-S12).

### The mathematical model

The model assumes a cell that is growing exponentially at the highest growth rate possible given a specific constant environmental setting. Thus, for each macromolecule in the cell (*c*), the sum of the rates (*v*) of all synthesis processes minus the sum of the rates of all degradation processes are balanced by the cellular growth rate *µ*:

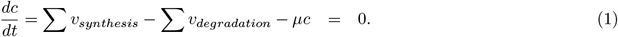

We solve for each cellular pool concentration (molecules per *µm*^3^ of biovolume) that maximizes *µ*. The reaction rates catalysed by a given protein *i*are represented by a Michaelis-Menten type equation:

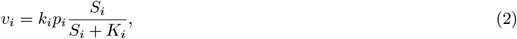

in which *k*_*i*_ is the maximum turnover rate per protein, *S*_*i*_ is the substrate concentration, *K*_*i*_ is the half-saturation constant, and *p*_*i*_ is the concentration of protein *i*in molecules of protein *i*per *µm*^3^ of biovolume. The glycolysis and lipid degradation fluxes estimated to meet dark respiration costs are an exception since all glucose and lipid storage will be used to fuel respiration during the night and thus Eq. 2 can be simplified as *v*_*i*_ = *k*_*i*_*p*_*i*_ for these pools.

We assume all enzymatic rates in the cell increase with temperature following the Arrhenius equation:

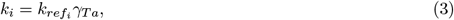

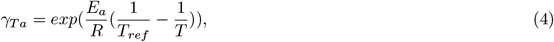

in which 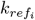 is the maximum turnover rate at a reference temperature *T*_*ref*_ (in Kelvin), *E*_*a*_ is the activation energy (eV), *R*is Boltzmann’s constant (eV K^−1^) and *T* is the actual temperature (K). The only exception is the maximum rate of photosynthesis (*k*_*p*_), which we assume is independent of temperature (*20*). Simulations were run where *E*_*a*_ was constant for all reactions (except photosynthesis) and where *E*_*a*_ varied for different rates (e.g. Figure 1B).

All protein pools *p*_*i*_ are synthesized by ribosomes (*p*_*ri*_), including ribosomes themselves. The total protein synthesis flux catalysed by ribosomes *v*_*ri*_ (molecules of amino acids *µm*^−3^ *min*^−1^) is partitioned among the different proteins according to the relative proteome investments *ϕ*_*i*_. Since our model does not simulate the full proteome, we assume that a fraction of the proteome devoted to other protein pools (*p*_*other*_; Table S2) is constant, and thus the sum of all proteome relative investments is equal to 0.5:

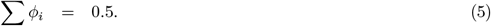

Our results are not sensitive to the choice of *p*_*other*_ (Figure S11). One can re-write Eq. 1 for a given protein pool *I*as following:

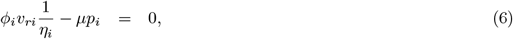

in which *η*_*i*_ is the molecular weight of a given protein (number of amino acids per protein). The model is fully defined in the Supplemental Material and detailed description of the different metabolic pathways are given in the following sections.

### Protein synthesis

The protein synthesis flux (*v*_*ri*_; molecules of amino acids *µm*^−3^ *min*^−1^) is catalysed by ribosomes and requires both internal carbon (*c*_*ic*_) and internal nitrogen (*c*_*in*_):

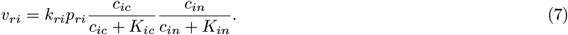

The cellular nitrogen to carbon demand 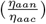is defined as:

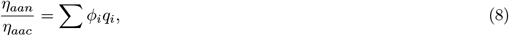

in which the subscript *i*represents protein pool *i*and *q*is the ratio of nitrogen to carbon for that protein pool. We use an average value for the carbon content of the protein pools (*η*_*aac*_, molecules of C/aa; Table S2) and solve for *η*_*aan*_ (molecules of N/aa). The above equation thus controls the consumption rate of carbon and nitrogen used in the synthesis of proteins to meet the optimal stoichiometry of the cell.

### Carbon metabolism

The conversion of dissolved inorganic carbon (DIC) into a generic pool of organic carbon (*c*_*ic*_) is mediated by the rubisco protein pool (*p*_*ru*_) such that the flux of carbon fixation *v*_*ru*_ (molecules of C *µm*^−3^ *min*^−1^) is given by:

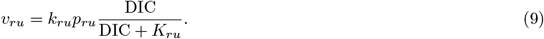

For the model presented in this paper, we assume that carbon is not limited: DIC *>> K*_*ru*_ so *v*_*ru*_ ≈*k*_*ru*_*p*_*ru*_. This internal organic carbon (*c*_*ic*_), in turn, is used in three different reactions: i) protein synthesis (*v*_*ri*_) catalysed by ribosomes; ii) lipid synthesis (*v*_*lb*_) catalysed by the protein pool *p*_*lb*_; and iii) glycolysis (*v*_*gl*_) catalysed by the protein pool *p*_*gl*_ to fuel cellular respiration during the night. The reaction rates *v*_*lb*_ and *v*_*gl*_ are in units of molecules of C *µm*^−3^ *min*^−1^.

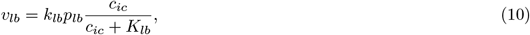

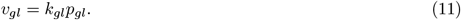

The glycolysis flux can be simplified as *v*_*gl*_ = *k*_*gl*_*p*_*gl*_ because we assume that all glucose will be used to fuel respiration during the night.

We can then write the equation for the internal pool of carbon:

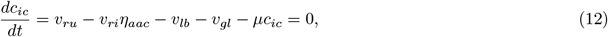

in which *v*_*ri*_ is multiplied by *η*_*aac*_ to convert the amino acid flux to carbon units.

### Nitrogen metabolism

Dissolved inorganic nitrogen (DIN) is imported into the cell (*v*_*tr*_; molecules of N *µm*^−3^ *min*^−1^) by nitrogen transporters (*p*_*tr*_) localized in the membrane:

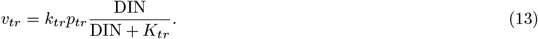

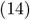

As we focus here on the impact of temperature on cell metabolism, we assume nitrogen is not limiting so that DIN*>> K*_*tr*_ and *v*_*tr*_ ≈ *k*_*tr*_*p*_*tr*_. Internal nitrogen (*c*_*in*_) is subsequently consumed by the protein synthesis pathway (*v*_*ri*_) such that the internal pool of nitrogen can be obtained from:

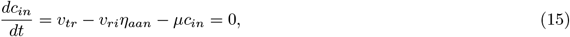

in which *v*_*ri*_ is multiplied by *η*_*aan*_ to convert the amino acid flux to nitrogen units.

### Cell stoichiometry

By simulating carbon and nitrogen metabolisms and varying the nitrogen content per amino acid (*η*_*aan*_) according to the nitrogen to carbon demand, it is possible to estimate the overall N:C quota of the photosynthetic cell (*q*_*cell*_) by computing the weighted average:

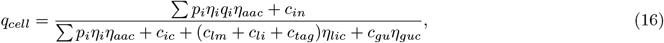

in which *i*represents protein pools, *p*is protein concentration, *η*is the protein molecular weight in units of aa protein^−1^, and *q*is the protein nitrogen to carbon ratio (see Table S2). It is important to note that *q*_*cell*_ represents the light proteome, thus accounting for the carbon storage (*c*_*gu*_ + *c*_*li*_) that is needed to meet dark respiration. As showed by previous studies, carbon storage required for dark metabolism can account for a significant amount of carbon in the cell (*15, 48*).

### Photosynthesis

Photosystems (*p*_*p*_) absorb photons at the rate of *v*_*p*_ (molecules of photon *µm*^−3^ *min*^−1^) converting light (*I*) energy into chemical energy at the rate of *v*_*ep*_ (molecules of energy *µm*^−3^ *min*^−1^), as follows:

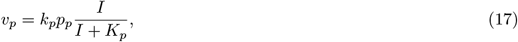

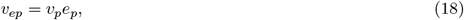

in which *e*_*p*_ is the amount of chemical energy produced per photon absorbed.

Photosystems are assumed to be damaged by heat, which causes photosystems to be dysfunctional in the model. It is known that photosystems are susceptible to heat damage, more specifically the D1 protein within photosystem II, causing its deactivation (*22*). We represent this process in the model as a temperature dependent damage term after Mairet *et al*. (*23*):

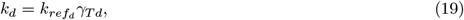

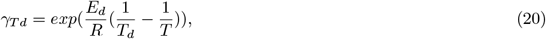

in which *k*_*d*_ is the maximum rate of damage at a given temperature, 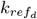 is the maximum rate of damage at a reference temperature *T*_*ref*_, and *E*_*d*_ is the deactivation energy which sets the slope at which temperature increases the damage of photosystems above *T*_*d*_.

We allow the cell to store lipids (*c*_*tag*_) which act to reduce the rate of photosystem damage. Lipid droplets act as a sink for reactive oxygen species and have been observed to play a role in the repair of damaged photosystems (*18*). The cell can thus choose to maximize the concentration of stored lipids up to 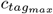 in the cell to minimize the rate of damage of photosystems as temperature changes, or it can choose to store less lipids at the cost of higher damage (Eqs. 21 and 22). Thus, the actual rate of damage *v*_*d*_ depends on *k*_*d*_, on the concentration of photosystems *p*_*p*_, and on the concentration of stored lipids *c*_*tag*_:

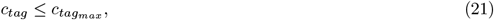

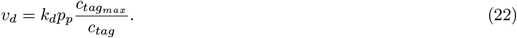

We assume that damaged photosystems (*p*_*dp*_) are repaired via repair proteins (*p*_*re*_) and that the rate of repair (*v*_*re*_) is calculated as:

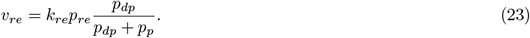

Considering that the repair processes are relatively fast compared to protein synthesis (*54*), the damage-repair processes can be assumed to converge to a quasi-steady state (*v*_*re*_ = *v*_*d*_), following Mairet *et al*. (*23*). This allows us to solve for the fraction of damaged photosystems:

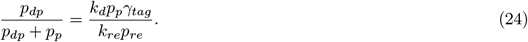

Finally, the photosystems equation can be written as:

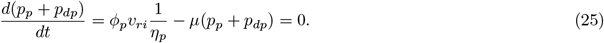

### Lipid metabolism

Lipids play three different roles in our model. First, we assume that the cell must invest in lipids to maintain membrane integrity following Molenaar *et al*. (*47*). This is implemented by assuming that the ratio between transporter proteins (*p*_*tr*_) and lipids (*c*_*lm*_) in the membrane must be equal to or lower than a certain value *M*_*min*_ + *M*:

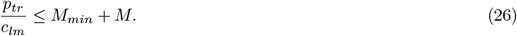

*M*_*min*_ is the minimum ratio between *p*_*tr*_ and *c*_*lm*_. The *M* term captures the temperature dependence of lipid saturation. Specifically, the saturation state of lipids increases as a function of temperature (*17, 18*). While unsaturated fatty acids have kinks in their molecular structure due to single or multiple double bonds, saturated fatty acids lack these kinks and thus can be more tightly packed together, occupying less space in the membrane. Since the constraint given in Eq. 26 controls the minimum spacing between transporters needed for membrane integrity, as temperatures increase, more lipids per surface area of membrane is needed. For simplicity, we assume that *M*decreases linearly as a function of temperature (*21*):

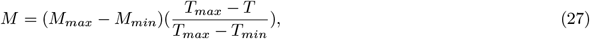

in which *M*_*max*_ is the maximum transporter to lipid ratio in the membrane and *T*_*max*_ and *T*_*min*_ are the maximum and minimum temperatures, respectively, at which we perform our modeling experiments.

Second, we assume that stored lipids *c*_*tag*_ can be used to mitigate oxidative stress as described above (Eqs. 21 and 22). Third, lipids can fuel dark respiration through the lipid degradation pathway. We thus optimize for the fractions of the lipid synthesis flux (*v*_*lb*_) that are used to synthesize: i) membrane lipids (*∝*_*lm*_), ii) stored lipids that can mitigate damage (*∝*_*tag*_) and iii) lipids for dark respiration (*∝*_*ld*_), so that:

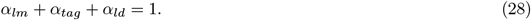

We assume that the lipid degradation pathway must be catalysed by the protein pool *p*_*ld*_, so that *v*_*ld*_ = *k*_*ld*_*p*_*ld*_ = *∝*_*ld*_*v*_*lb*_. Finally, we can define the equations for the membrane lipid and stored lipids pool as follows:

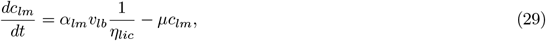

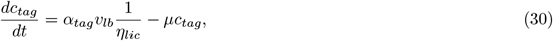

in which *v*_*lb*_ (molecules of carbon *µm*^−3^ *min*^−1^) is multiplied by the inverse of *η*_*lic*_ (molecules of C/lipid) to convert the flux from carbon to lipid units.

### Energy metabolism

The energy metabolism of the cell is constrained based on its energetic gains and costs, so that for each reaction rate *v*_*i*_, there will be an associated energy conversion factor *e*_*i*_ (Table S2). The cell can obtain energy through three pathways: photochemistry, glycolysis and lipid degradation. All other metabolic functions have an associated energetic cost. Energy fluxes are described in units of molecules of energy *µm*^−3^ min^−1^.

We first assume that light energy converted by photosystems must meet the energy cost associated with rubisco activity:

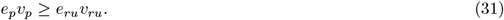

After meeting rubisco costs, photosystems can also contribute to meet the energetic demands of other metabolic functions (*e*_*cost*_):

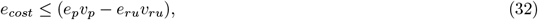

in which the total energetic cost of the cell (*e*_*cost*_), excluding rubisco costs, can be defined as the sum of the metabolic fluxes *v*_*j*_ multiplied by its respective energy costs *e*_*j*_:

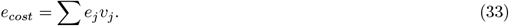

We assume that a fraction *f*_*dr*_ of *e*_*cost*_ happens in the dark and thus must be paid through either glycolysis (*v*_*gl*_) or lipid degradation (*v*_*ld*_):

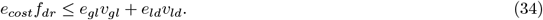

*f*_*dr*_ is a constant in our model and we assume it to be 25% of *e*_*cost*_. The overall metabolism of phytoplankton is slower in the dark and so *f*_*dr*_ can be expected to be lower than 0.5, but since this value is not well constrained we performed a sensitivity analysis to determine *f*_*dr*_ (Figure S12). Increasing *f*_*dr*_ shifts growth rates down and requires higher investment in respiratory pathways, but does not fundamentally change our conclusions (Figure S12). We can then calculate the concentration of the storage pools for both glucose (*c*_*gu*_) and lipids (*c*_*li*_) needed to fuel respiration during the dark period:

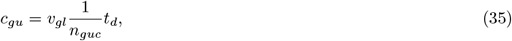

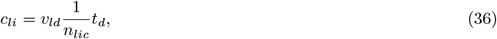

in which *n*_*gu*_ is the number of molecules of carbon per molecule of glucose, *n*_*li*_ is the number of molecules of carbon per molecule of stored lipid, and *t*_*d*_ is the duration in minutes of the dark period that the cell experiences over a day. Here, we assumed a 12:12 hours light-dark cycle.

### Space and density constraints

Following Molenaar *et al*. (*47*), we assume that the volume of the cell (*V*_*tot*_) is proportional to the total surface area (SA) occupied by transporters and lipids in the membrane multiplied by a factor *β*that corresponds to the volume to surface area ratio of the cell:

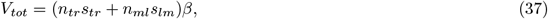

in which *n*_*tr*_ and *n*_*ml*_ are the total number of transporters and lipids in the membrane, and *s*_*tr*_ and *s*_*lm*_ correspond to the specific surface area of transporters and lipids molecules, respectively. The number of molecules of a given intracellular pool *i*can be linked to their concentrations as follows:

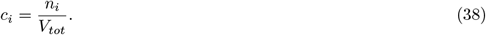

Thus, we can combine the equations above to get:

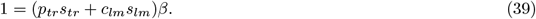

Unsaturated lipids are expected to have higher *s*_*lm*_ relative to saturated lipids due to the presence of kinks in their molecular structure. Previous studies found that *s*_*lm*_ can be up to 1.25 times higher for unsaturated lipids versus saturated lipids (*55, 56*). We performed sensitivity analyses assuming that *s*_*lm*_ decreases linearly with increases in temperature (due to increases in the saturation level of lipids; Eqs. 26 and 27) and found this to not impact our main results (Figure S13). Thus, to minimize free parameters in the model, we decided to assume a constant value for *s*_*lm*_.

Now we have only one unknown parameter *β* which can be optimized for. Since *β*is the volume to surface ratio of the cell, and assuming a spherical cell, we find that the cell radius (*r*) is equivalent to three times *β*:

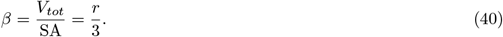

Note that other cellular geometries can be used with this model but for simplicity we use a sphere.

In order for the cell to maintain osmoregularity, we define a constraint on the maximum density of the cell. Photosystems are contained within the chloroplast and lipid droplets are hydrophobic and thus neither have a major effect on osmoregulation, but take up space. Thus, the maximum density of the cell (*D*_*max*_; Da *µm*^−3^) is defined relative to the volume of the cell that is not occupied by photosystems and lipid storage (*V*). We can then write an expression for the total volume of the cell:

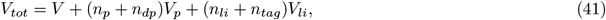

so that *n*_*p*_ and *n*_*dp*_ are the total number of functional and damaged photosystems and *n*_*li*_ and *n*_*tag*_ are the total number of stored lipids required to regulate oxidative stress and to fuel respiration, respectively. *V*_*p*_ and *V*_*li*_ correspond to the volume of photosystem and stored lipid molecules, respectively. We can re-write the expression above to obtain:

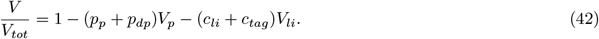

This expression allows us to define the concentration of the pools that affect the density of the cell relative to *V*within the density constraint:

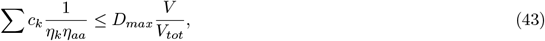

in which *k*is a subscript for all pools that influence the density of the cell, which includes all pools in the model except for photosystems, lipid storage and membrane components. *η*_*k*_ (number of amino acids per protein *k*) and *η*_*aa*_ (Da units per amino acid) converts *c*_*k*_ from units of molecule of protein per total cellular biovolume to Da units per total cellular biovolume (Table S2).

### Model validation and sensitivity runs

The model was compared against trait data obtained for several diatom species acclimated to different temperatures under controlled experimental conditions. The data were previously reported in Schaum *et al*. (*13*), Liang *et al*. (*24*) and O’Donnell *et al*. (*14, 35*), referred herein as the Schaum, Liang and O’Donnell datasets, respectively. Whenever needed, we used the free version of GetData Graph Digitizer http://getdata-graph-digitizer.com/ to digitize plots. Our modeling predictions were compared against the following traits: growth rates, carbon fixation rates, carbon use efficiency (i.e. 1 -respiration/photosynthesis), cell size (biovolume), cellular nitrogen to carbon ratio, and chlorophyll content per unit of biovolume.

The original source of the data used in Figures 2, S1, and 5 is as follows: (i) Schaum dataset: data in Figures 2 and S1 correspond to the control lineage evolved at 22°C, while data in Figure 5 also includes data for lineages evolved under moderate (26 °C) and severe (32 °C) warming. Growth rates, carbon fixation rates, and carbon use efficiency were reported in Figures 1 and 2 in Schaum *et al*. (*13*) while data for other traits (i.e. size, chlorophyll content and N:C ratio) were reported in Figure 3 in Schaum *et al*. (*13*); (ii) Liang dataset: we selected data from the warm-adapted strain CCMP160 grown at 24°C. Growth rates were given in Figure 1 while cell size and chlorophyll data were reported in Table 2 in Liang *et al*. (*24*); (iii) O’Donnell dataset: we validated our model against the cellular N:C ratios reported for two diatom strains grown at 16 °C and 31 °C in Figure 3 in O’Donnell *et al*. (*14*). To fit the model to the different growth curves, we tuned the parameters defining the thermal performance of metabolic reactions (i.e. 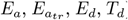) and the maximum rate of damage 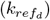. To fit for the cellular quota, we tuned the average N:C ratio of proteins (*q*_*pt*_). To fit the model to the observed cell sizes and chlorophyll content, we tuned the maximum cellular density (*D*_*max*_), the maximum ratio between transporters and lipids in the membrane (*M*_*max*_) and the maximum concentration of stored lipids 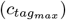. The latter directly impacts the energetic cost per photosystem damaged by heat-stress (*e*_*d*_). The values of the fitted parameters are given in Table S3.

We validated the model assuming that the cell could only use glycolysis (Figure 2) or lipid degradation (Figure S1) to fuel dark respiration, because our model solves for the optimal proteome of the cell in a steady-state environment and thus, results in a sharp transition between the two metabolic pathways (it is never optimal to use both pathways). The tuned parameters for the model runs in which the cell could only use the glycolysis pathway are given in Table S3. The validation of the model assuming that the cell could only use the lipid degradation pathway required just a few modifications to the values listed in Table S3, as follows: (i) Schaum: *D*_*max*_ = 1.0 × 10^10^ Da *µm*^−3^; (ii) Liang: *D*_*max*_ = 1.5 × 10^11^ Da *µm*^−3^ and *M*_*max*_ = 0.003; and (iii) O’Donnell: *D*_*max*_ = 1.6 × 10^11^ Da *µm*^− 3^.

Often conversion factors were necessary to compare the model output to experimental data. The simulated pool of photosystems was converted to units of chlorophyll per biovolume assuming that one photosystem has on average 140 molecules of chlorophyll (BNID 111790; Milo *et al*. (*57*) and considering that one molecule of chlorophyll has 893.5 g/mol. The carbon fixation rates reported in the Schaum dataset were converted from units of *µ*gC *µ*gC ^−1^ day ^−1^ to *µ*gC *µ*m ^−3^ day ^−1^ assuming a constant cellular carbon content of 1.25 × 10 ^−4^ pmolC *µ*m ^−3^ (Figure 3b in Schaum *et al*. (*13*)).

We solved the model in intervals of 1°C from 5°C to 45°C. We performed sensitivity analyses to investigate: i) how the thermal dependencies of membrane components influence cell size (Figure 4), ii) the impact of poorly constrained parameters on traits and proteome investments (Figures S11-S14) and iii) what influences the critical point at which the metabolic switch between glycolysis and lipid degradation occurs (Figure S7). The model was written in Julia (*58*) and the code will be deposited on GitHub and is currently available at https://dornsife.usc.edu/labs/levine/levinelabcode/.

## Acknowledgments

First and foremost, we are grateful for the valuable discussions with Dr. Noele Norris regarding model development and for her feedback on earlier versions of this manuscript. We would also like to acknowledge the comments by Dr. Ben Van Mooy when deciding how to model phytoplankton lipid metabolism and on all feedback received from Dr. Holly Moeller and members of the Levine Lab and the Moeller Lab in earlier versions of this manuscript.

## Author contributions

SGL and NML designed research. SGL and NML contributed to the development of the model and SGL performed research. SGL and NML analysed the data and wrote the paper.

## Data and materials availability

The model code will be deposited on GitHub and is currently available at https://dornsife.usc.edu/labs/levine/levinelabcode/.

